# Cerebellar GABA change during visuomotor adaptation relates to adaptation performance and cerebellar network connectivity: A Magnetic Resonance Spectroscopic Imaging study

**DOI:** 10.1101/2021.12.19.473367

**Authors:** Caroline R Nettekoven, Leah Mitchell, William T Clarke, Uzay Emir, Jon Campbell, Heidi Johansen-Berg, Ned Jenkinson, Charlotte J Stagg

**Affiliations:** Wellcome Centre for Integrative Neuroimaging, FMRIB, Nuffield Department of Clinical Neurosciences, University of Oxford (OX3 9DU); Oxford Centre for Human Brain Activity, Wellcome Centre for Integrative Neuroimaging, Department of Psychiatry, University of Oxford (OX3 7JX); Medical Research Council Brain Network Dynamics Unit, Nuffield Department of Clinical Neurosciences, University of Oxford (OX1 3TH); Department of Psychiatry, University of Oxford (OX3 7JX); School of Health Sciences, Purdue University (IN 47907); School of Sport, Exercise and Rehabilitation Sciences, University of Birmingham (B15 2TT)

**Keywords:** cerebellum, GABA, visuomotor adaptation, spectroscopy, neuroimaging, functional connectivity

## Abstract

Motor adaptation is crucial for performing accurate movements in a changing environment and relies on the cerebellum. Although cerebellar involvement has been well characterized, the neurochemical changes in the cerebellum underpinning human motor adaptation remain unknown.

We used a novel Magnetic Resonance Spectroscopic Imaging (MRSI) technique to measure changes in the inhibitory neurotransmitter γ-aminobutyric acid (GABA) in the human cerebellum during visuomotor adaptation. Participants used their right hand to adapt to a rotated cursor in the scanner, compared with a control task requiring no adaptation. We spatially resolved adaptation-driven GABA changes at the cerebellar nuclei and cerebellar cortex in the left and the right cerebellar hemisphere independently and found that simple right-hand movements increase GABA in the right cerebellar nuclei and decreases GABA in the left. When isolating adaptation-driven GABA changes, we found an increase in GABA in the left cerebellar nuclei and decrease in GABA in the right cerebellar nuclei. Early adaptation-driven GABA change in the right cerebellar nuclei correlated with adaptation performance: Participants showing greater GABA decrease adapted better, suggesting early GABA change is behaviourally relevant. Early GABA change also correlated with functional connectivity change in a cerebellar network: Participants showing greater decreases in GABA showed greater strength increases in cerebellar network connectivity. Results were specific to GABA, to adaptation and to the cerebellar network.

This study provides first evidence for plastic changes in cerebellar neurochemistry during motor adaptation. Characterising these naturally occurring neurochemical changes may provide a basis for developing therapeutic interventions to facilitate human motor adaptation.

**Significance Statement:** Despite motor adaptation being fundamental to maintaining accurate movements, its neurochemical basis remains poorly understood, perhaps because measuring neurochemicals in the human cerebellum is technically challenging. Using a novel magnetic resonance spectroscopic imaging method, this study provides evidence for GABA changes in the left and right cerebellar nuclei driven by both simple movement and motor adaptation. The adaptation-driven GABA change in the right cerebellar nuclei correlated with adaptation performance, and with functional connectivity change in a cerebellar network. These results show the first evidence for plastic changes in cerebellar neurochemistry during a cerebellar learning task. This provides the basis for developing therapeutic interventions that facilitate these naturally occurring changes to amplify cerebellar-dependent learning.

## Introduction

As we move through the world we are constantly confronted with changes in our bodies and our environment. Muscle properties change due to fatigue, exercise or development and the objects we interact with vary in weight, size or how well we can manipulate them. Adapting our motor commands to these variations is thus crucial to interacting with our environment.

The cerebellum plays a key role in motor adaptation. Patient studies and brain stimulation suggest that an intact cerebellum function is a prerequisite for adaptation (Martin et al., 1996; Tseng et al., 2007; Taylor et al., 2010; Diedrichsen et al. 2005, Jenkinson & Miall, 2010), with more severe cerebellar impairment resulting in worse adaptation (Tseng et al., 2007). Healthy participants show increased cerebellar activity early in adaptation (Seidler et al., 2006; Ichise et al., 2000) and in response to sensory prediction errors (Schlerf et al., 2012), as well as adaptation-related changes in cerebellar activity in visuomotor learning paradigms (Miall and Jenkinson 2005; Graydon et al., 2005). Though these studies provide a compelling argument for cerebellar involvement in adaptation, the physiological changes in the cerebellum during adaptation are not yet fully understood.

Adaption may lead to decreases in the major inhibitory neurotransmitter γ-aminobutyric acid (GABA). GABA is found throughout the cerebellum and plays an important role in cerebellar plasticity (Ito, 2014). In the cerebellar cortex, GABA has been implicated in plasticity at parallel fibre synapses (Orts-Del’Immagine et al., 2018) as well as Purkinje cell synapses (He et al., 2015). GABAergic Purkinje cells, the sole output neurons of the cerebellar cortex, exert a constant inhibitory tone on the deep cerebellar nuclei via frequent simple spike firing. Strong climbing fibre input causes Purkinje cells to fire complex spikes, resulting in a pause in simple spike firing, hence releasing the deep cerebellar nuclei from inhibition (Thach, 1967). In nonhuman primates, Purkinje cells increased complex spike firing, and therefore decreased inhibition, after introduction of a perturbation, which returned to baseline as the monkeys learned to adapt their movements and errors decreased (Gilbert et al., 1977). However, despite the importance of GABA in motor adaptation, cerebellar GABA changes during human motor adaptation have not yet been investigated, likely due to the technical challenges presented by cerebellar MR in dealing with the low signal-to-noise ratio in the cerebellum, which is located deep in the cranium and close to brain stem structures that move during the cardiac cycle (Brooks et al., 2013; Diedrichsen et al., 2010).

In this within-subject, crossover study, we used a novel Magnetic Resonance Spectroscopic Imaging (MRSI) approach to quantify GABA in the human cerebellum while participants performed a visuomotor adaptation task with their right hand, compared with a non-rotated control using the same hand. We were able to spatially resolve adaptation-driven GABA changes at the cerebellar nuclei and in the cerebellar cortex in the left and the right cerebellar hemisphere independently. We hypothesised that adaptation would lead to a decrease in GABA in the right deep cerebellar nuclei, as the uncrossed pathways at the level of the cerebellum results in mainly ipsilateral hand representations (Buckner et al. 2011, Mottolese et al. 2013, Diedrichsen and Zotow, 2015). We hypothesised that the degree of GABA decrease would be correlated with adaptation on a subject-by-subject basis, such that greater GABA decreases related to greater adaptation. Finally, since previous studies have shown an increase in intrinsic functional connectivity in the cerebellum after adaptation (Albert et al., 2009; Nettekoven et al., 2020) and functional connectivity has been shown to correlate with GABA in a major network node (Stagg, 2014, Bachtiar 2018), we hypothesised that GABA decrease would correlate with increased cerebellar functional connectivity.

## Materials and Methods

### Participants

Seventeen healthy, right-handed participants (6 female; 18-34 years; mean age 23 years) participated in this within-subject, crossover study. Each participant attended two sessions during which they performed a visuomotor task with either an adaptation component (rotation condition) or no adaptation component (control condition) during MRI. The order of the sessions was counterbalanced across the group. Participants had no history of neurological or psychiatric conditions and were not taking any psychoactive medications. All participants gave their written informed consent to participate, in line with central university research ethics committee approval (University of Oxford; R59564/RE002).

### Experimental protocol

The experimental protocol for each session is shown in figure 1A. Participants performed a visuomotor task using a hand-held joystick (Nata Technologies, custom-built fMRI joystick, 60 Hz sampling rate, figure 1C left panel) to control a cursor on a screen, which either involved an adaptation component (rotation condition) or no adaptation component (control condition). Participants were instructed to make ballistic movements to ‘shoot’ with their cursor through one of eight targets (Figure 1C middle panel).

**Fig. 1.**
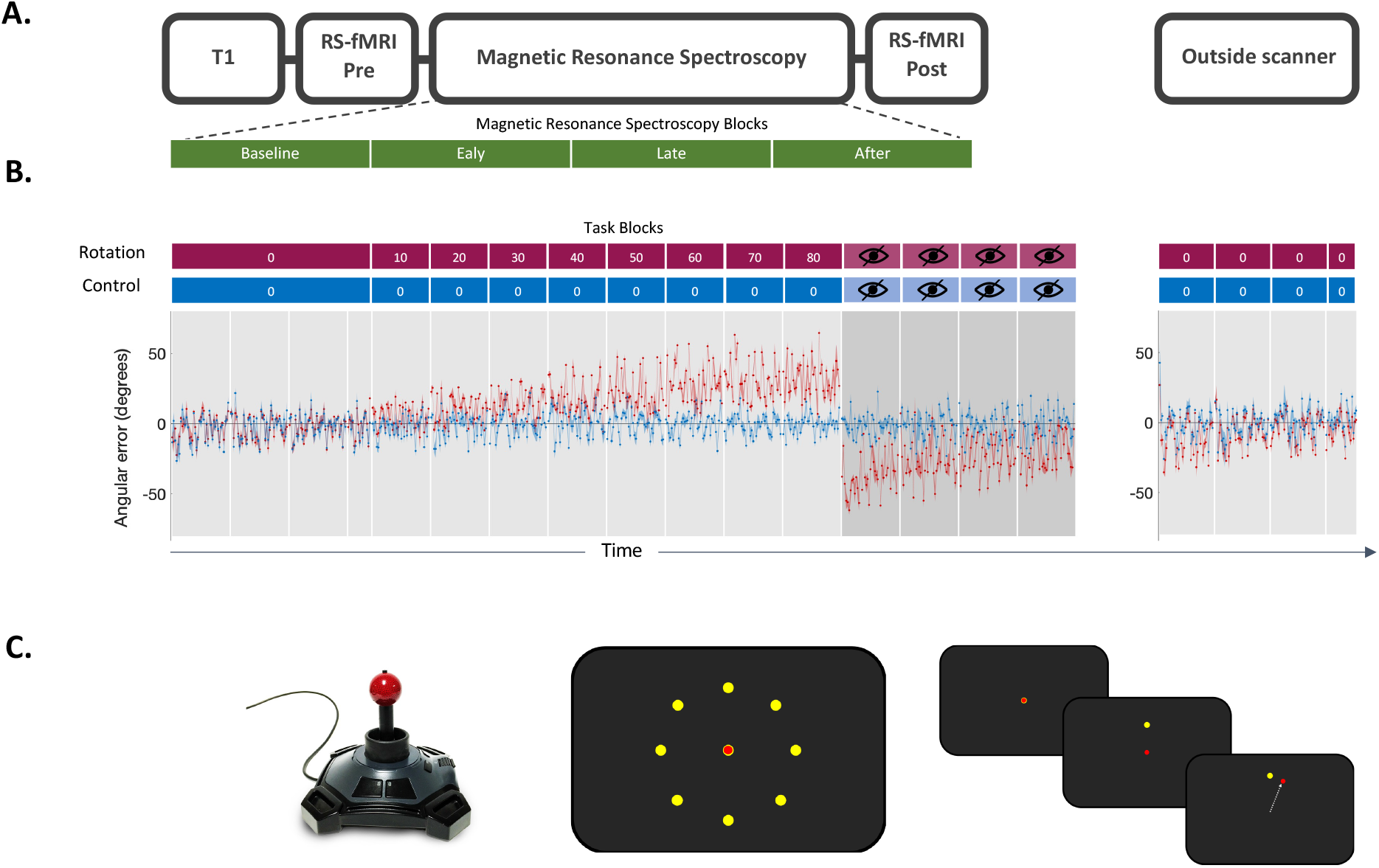
Experiment. **A.** Scanning Protocol. T1-weighted structural image was acquired at the begin of the MRI scan. Resting-state fMRI data was acquired before and after the task. Magnetic Resonance Spectroscopy Imaging (MRSI) data was acquired during the task. Four MRSI scans were acquired during performance of the visuomotor task with each MRSI scan lasting 9 min, total acquisition 36 min. **B.** Behavioural data. Participants used a joystick to shoot targets on a screen. Participants began by performing 136 trials with no rotation imposed serving as the baseline in both conditions. In the rotation condition (red blocks), stepwise increasing rotated visual feedback required participants to adapt movements to reduce errors. One block at each angle and each block consisted of 40 trials of 4 seconds duration each. The numbers in the red and blue boxes indicate the degree to which the visual feedback was rotated, with 0° indicating no rotation. The imposed rotation reached a maximum of 80°, after which visual feedback was removed for four blocks of 40 trials each (blocks with crossed out eye). In the control session (blue blocks), participants performed the task without any rotation imposed, but for the same length (480 trials in total for the main task). The rotation was washed out after task (144 trials, no rotation). The task was practised before the main task outside of the scanner (32 trials, no rotation; not depicted in this figure). Behavioural data is shown as angular error at each trial averaged across participants. Shaded area represents standard error of the mean. Rotation condition error is shown in red. Control condition error is shown in blue. **C.** Task Schematic. **Left.** MR-compatible Joystick used to record participant responses. **Middle.** Eight possible target locations (yellow) centred radially around the cursor (red) at its starting position. **Right.** Schematic of a rotation trial. Cursor (red) is first presented at the centre starting position. Target (yellow) appears at one of the eight possible target locations. Participant makes a centre-out movement towards the target, but sees clockwise rotated visual feedback.

The target appeared at one of eight locations radially aligned around the starting position of the cursor and separated by 45°. Trials were grouped into epochs, with one epoch containing eight consecutive trials and all blocks containing five epochs (total: 5×8 = 40 trials; one block equates to one grey shaded region in figure 1B). The sequence of target presentation was fixed across participants. Participants had to perform the movement within a time window of 750 ms, after which the target disappeared, to ensure short, ballistic movements, minimising online corrections. No endpoint feedback was provided.

At the beginning of each session, participants performed a familiarisation block outside the scanner which consisted of 40 trials with no rotation imposed. They then performed 136 trials of no rotation in the scanner, serving as the baseline. Then, in the rotation condition, participants were required to adapt their centrifugal shooting movements to a rotation of the visual feedback which increased stepwise by 10 degrees after every block of 40 trials to maximally drive adaptation throughout the duration of the scanning session (Figure 1B red blocks). The imposed rotation reached a maximum of 80°, after which visual feedback was removed for four blocks of 40 trials each (blocks with crossed out eye in figure 1B) to assess aftereffect. In the control condition, participants performed the task without any rotation imposed for the full duration of the task (Figure 1B blue blocks). After the scan, participants performed 144 trials (i.e. 3.6 blocks) with no rotation imposed to probe retention of the previously learned compensatory movement.

### Behavioural analysis

Cursor movements were analyzed on a trial-by-trial basis using in-house software written in Matlab (Mathworks Inc, Natick, USA). The joystick position data (X and Y) was collected at a sample rate of 60 Hz. The kinematic data was filtered with a zero-phase filter with a 25 Hz cut-off and numerically differentiated to determine velocity. Trials that showed premeditated or incomplete movements were excluded from further analysis (mean number of rejected trials 30 ± 29).

For each trial, the angular error (measured in degrees) was calculated as the angle between a line connecting the starting position with the position of peak velocity of the cursor and the line connecting the starting position with the target. Positive values indicate a clockwise error (‘overshooting’) and negative values indicate a counter-clockwise error (‘undershooting’).

We quantified adaptation in the rotation condition by calculating the mean error across all rotation blocks, excluding the first epoch of each block (Galea et al., 2011). We normalised the mean error in the rotation condition to the mean error in the control condition to ensure that differences in the adaptation measure were specific to better adaptation to the imposed rotation. To do this, we calculated error change in the rotation condition by dividing the angular error in rotation blocks by angular error at baseline. We then calculated the difference between error change in the rotation condition and error change in the control condition by subtracting control from rotation error change. Positive change values therefore reflect a greater increase in error in the rotation condition than in the control condition. The normalisation formula applied to the rotation condition error was the same as the GABA normalisation, which is described in a later section. We refer to this normalised rotation error as adaptation error.

To quantify aftereffect, the mean error during all open loop trials of the rotation condition was normalised to the mean error during the control condition open loop trials. We refer to this normalised open loop measure as aftereffect. To quantify retention, the mean error in the first washout block of the rotation condition was normalised to the mean error in the first washout block after the control condition. We refer to this normalised washout error as retention error.

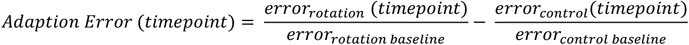

To determine whether participants adapted in the rotation condition, we used the R package lme4 (Bates et al. 2015) to construct a linear mixed effects model (LME) of error with epoch and target as fixed effects. To directly compare error reduction from the beginning of each block to the end of each block in the rotation condition and in the control condition, we constructed an LME model of the first and last epoch error in both conditions and tested for a significant condition x epoch interaction. In this LME model, we also entered target as a fixed effect to account for target-specific movement biases. For all LME models we allowed intercepts for different subjects to vary, to account for covarying residuals within subjects.

### Neuroimaging acquisition

All Magnetic Resonance (MR) data were acquired on a 3T Siemens scanner (PRISMA) with a 32-channel head coil. Structural MRI data were acquired using a magnetisation prepared rapid acquisition gradient echo (MPRAGE) sequence (Repetition Time 1900 ms, TE 3.96 ms, slice thickness 1.0 mm, in-plane resolution 1.0 x 1.0 mm^2^, flip angle = 8, FOV 256 x 232 mm^2^). The resting-state fMRI acquisition was matched to the UK Biobank resting-state fMRI parameters described in (Alfaro-Almagro et al. 2018). A multiband isotropic EPI acquisition (TR/TE = 735/39 ms, FOV = 210 x 210 mm^2^, bandwidth = 2030 Hz/Px, multiband acceleration factor 8, voxel dimension = 2.4 x 2.4 x 2.4 mm^3^, whole brain, acquisition time = 6:10 min for a total of 490 volumes) was performed before and immediately after the task. A separate “single-band reference scan” (SBRef) was acquired before the multiband acquisition (Moeller et al. 2010). This serves as the reference scan during pre-processing (e.g. during motion correction and registration steps), as it suffers from the same distortions as the multiband data, but has higher contrast (Alfaro-Almagro et al. 2018). Subjects fixated on a cross-hair image presented centrally on the screen during resting-state fMRI acquisition.

### fMRI analysis

All MRI data processing was carried out using FSL (FMRIB’s Software Library, Version 6.00, Jenkinson et al. 2012). For resting-state functional data, pre-processing steps were largely matched to the UK Biobank processing pipeline (Alfaro-Almagro et al. 2018). The data was motion corrected using MCFLIRT (Jenkinson et al. 2012), slice-timing corrected using Fourier-space time-series phase-shifting, distortion-corrected using fieldmaps (B0 unwarping), stripped of non-brain voxels using BET (Smith 2002), grand-mean intensity normalised and highpass temporally filtered (Gaussian-weighted least-squares straight line fitting, with sigma = 50.0 s)).

To remove structured noise, functional data were cleaned using a single-subject ICA, implemented in MELODIC and the automated classifier FIX, which was trained on 40 hand-labelled Biobank datasets to identify noise ICA components (Beckmann et al., 2004; Salimi-Khorshidi et al. 2014; Griffanti 2014). Although these training datasets were matched in acquisition sequence and experimental parameters to the datasets acquired in our experiment, we checked its classification accuracy to ensure the suitability of the Biobank-trained FIX algorithm for our data by manually classifying four datasets and comparing our labels to the FIX-classification. The FIX-classified labels matched our hand classification at 99.8%, which is within the expected range of classification accuracy (Alfaro-Almagro et al. 2018).

Functional data were registered to the MNI-152 template, using the SBref image as an intermediate step. Functional data were first linearly registered to the SBref image using 6 degrees of freedom (FLIRT; Jenkinson et al. 2001; Jenkinson et al. 2002), then to the structural image using boundary-based registration (BBR; implemented in FLIRT), and finally to the MNI-152 template using non-linear registration (FNIRT; Andersson et al., 2007; Andersson et al., 2019). Finally, spatial smoothing was applied with a 5 mm FWHM Gaussian kernel.

### Group Networks

In order to derive group networks, we applied a group ICA. For this, we concatenated the pre-processed functional data temporally across sessions and across subjects to create a single dataset. We then used the group-ICA to obtain 50 components. Visual inspection of the group-level components revealed that at a dimensionality of 50 the cerebellar network of interest included only cerebellar regions, whereas at a lower dimensionality, this network included regions of the visual cortex. We confirmed the cerebellar network (Figure 5A) as the network with the greatest spatial correlation with cerebellar regions active during performance of sensorimotor tasks (Hardwick et al., 2013). To determine the anatomical specificity of any effects we identified the Default Mode Network (DMN; Figure 5B) as a control network which we expected to be unaffected by task condition, and which does not spatially overlap with the task activation mask.

### Dual regression

Subject-and-scan-specific networks were obtained via a dual regression approach (Nickerson 2017), masked by the group mean network. We quantified network strength for the network of interest by extracting the mean z-value from the subject-and-scan-specific network maps masked by the group network mask.

### Connectivity change

We tested whether change in network strength correlated with change in GABA. To do this, we calculated change in network strength using the same normalisation procedure as the one used for behavioural data, namely dividing the strength value of the subject’s network after task performance (post; Figure 1A) by network strength at baseline (pre; Figure 1A) and calculating the difference between rotation strength change and control strength. Positive change values therefore reflect a greater increase in network strength in the rotation condition than in the control condition. We refer to this measure as connectivity change.

### Spectroscopic Imaging acquisition

Structural images were used to manually place a 65 x 25 x 15 mm^3^ MRSI slab in the cerebellum. The slab was positioned to cover anterior and superior areas of the right cerebellum which contain the hand motor representation (Buckner et al. 2011, Mottolese et al. 2013, Diedrichsen and Zotow, 2015), while avoiding contact with either the dura, to minimize significant macromolecule contamination, or with brainstem, to avoid excess physiological noise (Brooks et al., 2013).

Four MRSI scans were acquired during performance of the visuomotor task (time per scan = 9 min, total acquisition 36 min, TE 32 ms; Figure 1A). MRSI data was acquired using a nonwater suppressed semi-LASER DW-CRT sequence with metabolite cycling (Steel et al. 2018) at 5 x 5 x 15 mm^3^ resolution.

### Spectroscopy analysis

MRSI data was reconstructed and pre-processed according to Steel et al., 2018 using in-house scripts. Briefly, pre-processing included: metabolite cycling reconstruction (Emir et al 2017), coil-combination (Walsh et al., 2000), and correction for frequency and phase shifts, residual water removal using HLSVD (Cabanes et al., 2001) and eddy currents correction using the unsuppressed water signal (Klose, 1990). Concentration of neurochemicals was quantified as in Steel et al., 2018, using LCModel (Provencher et al., 2001). Specifically, we used a chemical shift of 0.5 to 4.2 ppm, a basis set containing 30 metabolites, default LCModel macromolecules, soft constraints on metabolites were disabled (NRATIO set to 0), and a baseline stiffness setting (DKMTM) of 0.25 (raw spectrum, spectral fit and scaled basis spectra in figure 2C). Voxels in the metabolite maps were excluded if CRLB of the respective metabolite > 60%, LCModel defined linewidths (FWHM) > 20 Hz, or signal-to-noise ratio (SNR) < 15 (thresholded GABA shown maps in figure 2A). MRSI data analysis therefore yielded independent quantification of neurochemical concentrations at four timepoints. All neurochemical concentrations are expressed as a ratio to total Creatine (Cr + PCr, tCr).

**Fig. 2.**
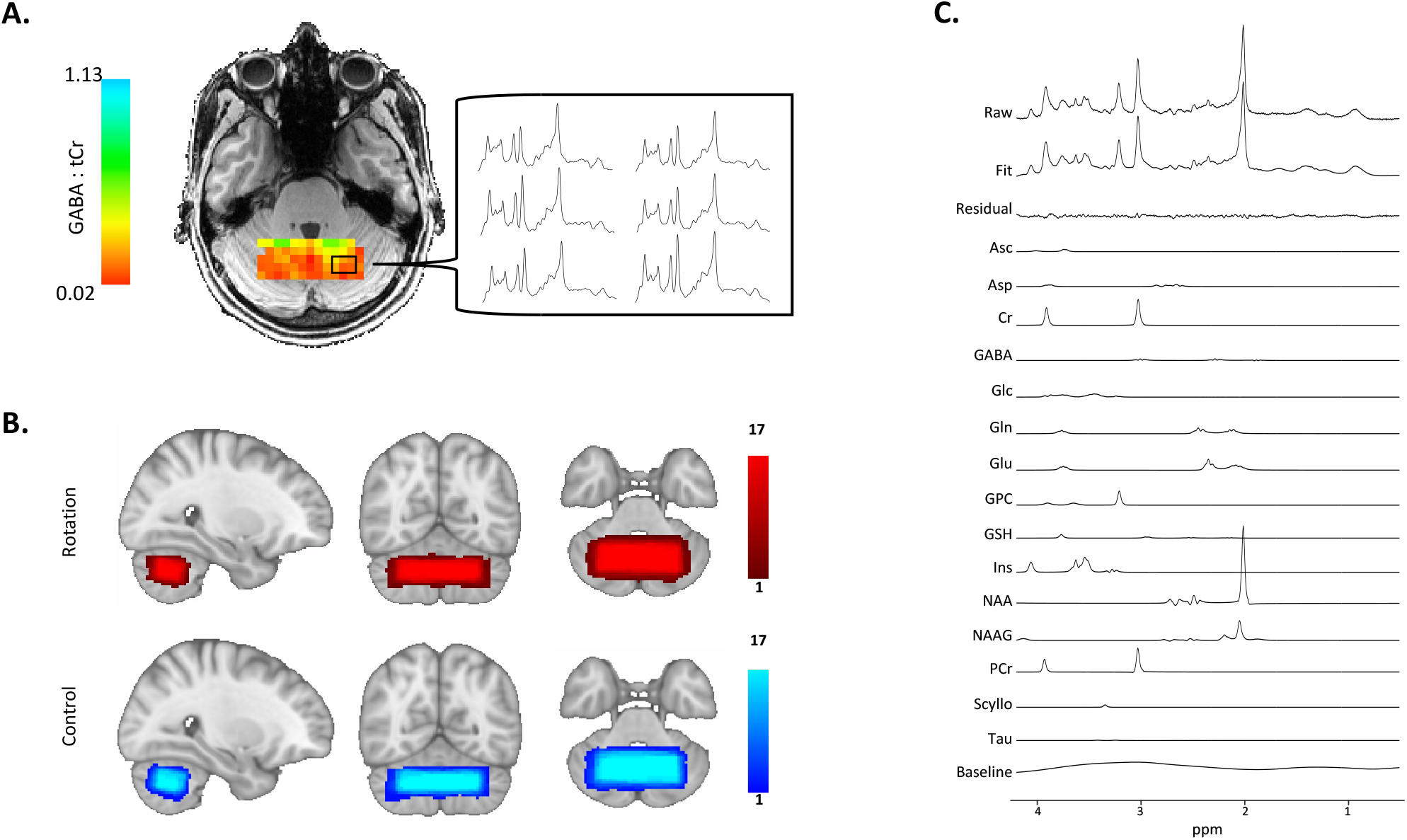
Cerebellar Magnetic Resonance Spectroscopic Imaging. **A.** Representative GABA Map. Image shows representative GABA map for one subject after quality control thresholding including the individual MRSI spectra from six voxels which allow quantification of GABA concentration in each voxel. Colour bars show GABA:tCr concentration in each voxel. **B.** Voxel Placement. MRSI voxel overlap maps for the two conditions. 65 × 25 × 15 mm3 MRSI slab placed over the anterior and superior areas of the right cerebellum such that it covers the hand motor representation. Colour bars represent number of participants. **C.** Spectral Fit. Image shows representative spectra of one subject including LCModel fit. Metabolite basis spectra have been scaled using LCModel such that a linear combination of the basis spectra, the residual and the baseline best fits the raw measured spectrum. Measured raw spectrum is shown in the at the top and LCModel Fit is shown below. GABA metabolite peaks appear at 1.89 ppm, 2.29 ppm and 3.01 ppm.

Metabolite maps were registered to the MNI template by a linear registration to the structural scan (FLIRT; Jenkinson et al., 2001; Jenkinson et al., 2002), and then non-linearly warp to the MNI template (FNIRT; Andersson et al., 2007; Andersson et al., 2019). Overlap maps of the MRSI slab in MNI space for each condition are shown in figure 2B. The MRSI slab was reproducibly placed between sessions (mean spatial correlation coefficient between each subject’s pair of MRS voxels: r = 0.85 ± 0.06).

To address our hypotheses, we conducted an ROI-based analysis using the SUIT toolbox (Diedrichsen, 2006) for SPM12. We created SUIT-space ROIs for the left and right cerebellar nuclei and cortex separately (Figure 3C right panel and Figure 3I right panel). We then registered the SUIT-space ROI masks to each subject’s structural scan.

**Fig. 3.**
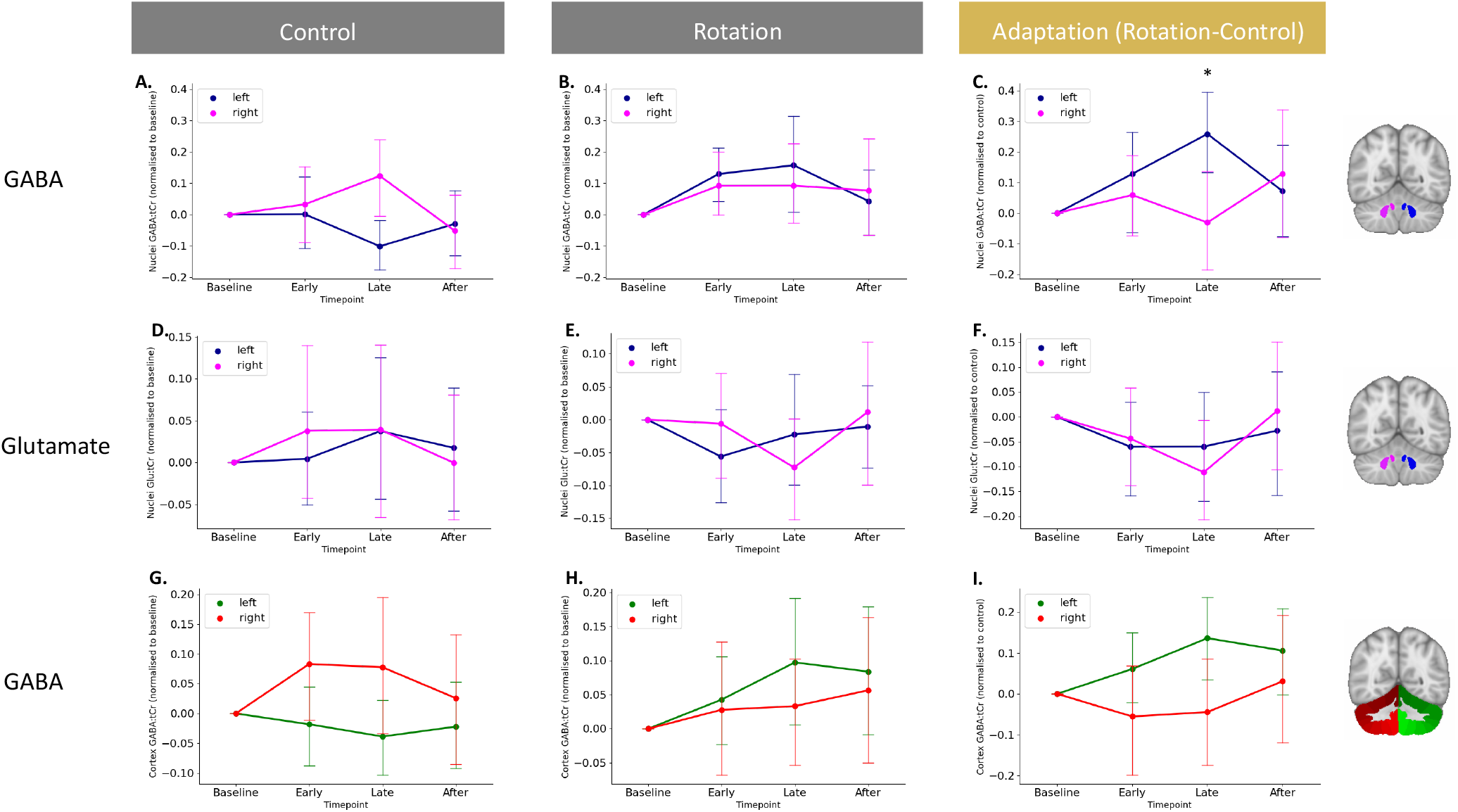
Isolating adaptation reveals GABA diverges in the left and right cerebellar nuclei. **A.** GABA changes in the cerebellar nuclei during movement execution. Shown are GABA values normalised to baseline in the control condition. **B.** No GABA changes in in the left and right cerebellar nuclei while participants in the rotation condition. Shown are GABA values normalised to baseline in the rotation condition. **C.** Adaptation-driven GABA changes in the left and right cerebellar nuclei. Against a backdrop of GABA changes related to simple movement execution, GABA diverges in the left and right cerebellar nuclei driven by adaptation. **D-F.** Glutamate does not change in the left and right cerebellar nuclei during the control condition (**D**), rotation condition (**E**) or when isolating adaptation (**F**). **G-I.** GABA does not change in the left and right cerebellar cortex during the control condition (**G**), rotation condition (**H**) or when isolating adaptation (**I**). For visualisation purposes, data in left and middle column is shown normalised to baseline. Statistics for control and rotation data was conducted on raw, non-normalised data. Statistics testing for adaptation-driven changes were calculated on rotation data normalised to control data.

Normalisation.

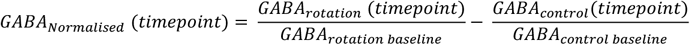

To control for movement effects, we normalised GABA in the rotation condition to GABA in the control condition using the same normalisation procedure as was applied to the behavioural data. The normalised GABA value at each timepoint is therefore equivalent to taking the difference between the percentage change in rotation GABA and the percentage change in control GABA. The GABA data was normalised for each hemisphere separately, resulting in normalised values for left and right cerebellar hemisphere (GABA_Left_ and GABA_Right_ respectively).

### Statistical Analysis

Mean ± SD are presented throughout. Statistical analyses of the data were conducted using R (RCoreTeam2013) and Statistics Package for the Social Sciences (SPSS, Version 25, IBM Corp., Armork, NY, USA). Repeated Measures-ANOVAs were used to test for neurochemical changes in the cerebellum during the different experimental conditions in the left compared to the right cerebellum. Maunchly’s test of sphericity was used to test the homogeneity of variance. Where Mauchly’s test of sphericity was significant (p<0.05) in RM-ANOVAs, Greenhouse-Geisser corrections were applied. Post hoc tests were calculated using Bonferroni-adjusted paired-samples t-tests.

To test relationships between neurochemical changes and behaviour, Pearson’s correlation coefficients (two-tailed) were calculated. Partial correlations were used to control for a behavioural index of no interest (two-tailed, unless otherwise specified). To determine whether two correlation coefficients were significantly different, Pearson’s correlation coefficients were converted into z-scores using Fisher’s Z-transformation. One-tailed z-tests were then performed on the z-scores given the standard error of each z-score (Myers et al., 2006).

## Results

### Participants adapted in the rotation condition and retained the aftereffect

We first tested whether participants adapted by reducing their error to the applied visuomotor perturbation in the rotation condition. As expected, we found that participants adapted within rotation blocks, indicated by a significant effect of epoch on error (LME of error in rotation blocks: χ2(1) = 5.78, p = 0.02; Figure 1B). To test whether this error reduction was specific to the rotation condition, we then compared error reduction in the rotation condition and error reduction in the control condition directly. In line with error reduction taking place exclusively in the rotation blocks of the rotation condition, we observed a significant interaction of epoch and condition in an LME (χ2(1) = 12.697, p < 0.001).

We next tested whether participants retained the adapted state. We observed an aftereffect in each of the four open loop blocks of the rotation condition, indicated by a significant negative error (Bonferroni-corrected one sample t-test of mean error for each open loop block, all p’s < 0.001). In contrast, no aftereffect was observed in any of the open loop blocks of the control condition (Bonferroni-corrected one sample t-test of mean error for each block, all p’s >0.99). Directly comparing the open loop blocks of the rotation condition and the control condition confirmed the presence of an aftereffect throughout the open loop blocks (Bonferroni-corrected paired t-test of mean error for each open loop block, all p’s < 0.001; Figure 1B).

We found that participants retained the compensatory movement in the washout phase of the rotation condition (Bonferroni-corrected one sample t-test of mean error for each block, all p’s < 0.01), but that there was no error during the washout phase of the control condition (Bonferroni-corrected one sample t-test of mean error for each block, all p’s > 0.28).

Because the rotation task imposed an adapted state that persisted throughout the washout phase, we tested for potential carry-over effects from the rotation session to the next session in this counterbalanced study. We found no evidence for carry-over effects in the second session, since there was no difference in baseline error of the second session between participants who had performed the rotation condition in the first session and participants who had performed the control condition in the first session (Paired t-test of baseline error: −4.090 ± 0.536, t(15) = −1.760, p = 0.099).

### GABA is stable across sessions

Because the MRSI approach employed in this study has not been used before in the cerebellum, we first quantified the reliability of our GABA measurements at baseline across sessions. The intraclass-correlation coefficient (ICC) of the baseline GABA measurements was 0.45 (F(16,16.5) = 2.63; p = 0.03), indicating a ‘fair’ reliability between sessions. To check whether GABA quantification was affected by the number of MRSI voxels included in an ROI, we looked for a relationship between GABA:tCr and number of voxels in our ROIs, and found no relationship in either the deep cerebellar nuclei masks (r = −0.13, p = 0.29) or the cerebellar cortex masks (r = −0.1 p = 0.42). Further, there was no difference in the number of voxels surviving quality assessment across the slab between the two conditions (t(16) = 1.140, p = 0.27).

### GABA changes in the cerebellar nuclei during movement execution

We first investigated GABA changes over time between conditions, regions, and hemispheres, via a four-way RM-ANOVA with within-subject factors of condition (Rotation, Control), region (Nuclei, Cortex), hemisphere (Left, Right) and timepoint (Base, Early, Late, After). There was no main effect on GABA:tCr of condition (F(1,16) = 1.145, p = 0.300, partial η2 =.067) or timepoint (F(3,48) = 1.391,p = 0.257, partial η2 =.08). However, GABA:tCr differed between regions (F(1,16) = 463.532, p < 0.0001, partial η2 =.967) and between hemispheres (F(1,16) = 71.816, p < 0.0001, partial η2 =.818), with higher GABA:tCr in the cerebellar nuclei (0.554 ± 0.014) than in the cerebellar cortex (0.348 ± 0.012) and higher GABA:tCr in left (0.478 ± 0.013) compared to the right cerebellar hemisphere (0.424 ± 0.012). There was a significant interaction between region and hemisphere (F(1,16) = 21.403, p < 0.001, partial η2 =0.572), driven by a greater difference in GABA:tCr between the hemispheres in the nuclei than in the cortex. Finally, we observed a significant three-way interaction between condition, hemisphere and timepoint (F(3,48) = 4.544, p = 0.007, partial η2 = 0.221). All other interactions were non-significant (all p > 0.1).

To understand this three way interaction, we performed RM-ANOVAs on GABA:tCr from the two conditions and the two regions separately, with within-subject factors of hemisphere (Left, Right) and timepoint (Base, Early, Late, After; Figure 3).

In the control condition, there were no bilateral task-related changes in GABA:tCr in the cerebellar nuclei (main effect of timepoint F(3,48) = 0.793,p = 0.504, partial η2 =0.047), but there was a significant main effect of hemisphere (F(1,16) = 9.551, p = 0.007, partial η2 =0.374) and a significant hemisphere*time interaction (F(3,48) = 3.943, p = 0.014, partial η2 =0.198; Figure 3A). Post hoc t-tests demonstrated that GABA:tCr was higher in the left than right nuclei at baseline (t(16.000) = 3.552, p = 0.011, 0.829 ± 0.963). However, during control task performance, GABA decreased in the left nuclei and increased in the right nuclei, so that there was no significant difference between the hemispheres during task performance (Early: t(16.000) = 2.204, p = 0.170, 0.636 ± 1.190; Late: t(16.000) = −0.438, p = 1.000, −0.125 ± 1.179; Figure 3A shows GABA:tCr timecourse normalised to baseline). In the cerebellar cortex, performance of the control task had no significant effects on GABA:tCr (main effect of hemisphere: F(1,16) = 3.166, p = 0.094, partial η2 =0.165; main effect of timepoint: F(3,48) = 0.793,p = 0.275, partial η2 =0.017; hemisphere*time interaction: F(3,48) = 3.943, p = 0.277, partial η2 =0.074; Figure 3G).

In the rotation condition, there were no task-related changes in GABA:tCr in the cerebellar nuclei (main effect of timepoint: F(3,48) = 2.528,p = 0.068, partial η2 =0.136; hemisphere*time interaction: F(3,48) = 0.539, p = 0.658, partial η2 =0.033; Figure 3B) or the cortex (main effect of timepoint: F(3,48) = 0.644,p = 0.591, partial η2 =0.039; hemisphere*time interaction: F(3,48) = 0.539, p = 0.658, partial η2 =0.033; Figure 3H).

### Isolating adaptation reveals GABA diverges in the left and right cerebellar nuclei

Given the significant effects of movement on GABA:tCr in the cerebellar nuclei, we wished to isolate any effects of adaptation over and above movement on inhibition. To do this we normalised GABA:tCr during the rotation condition to GABA:tCr during control by subtraction (Figure 3C and Figure 3I). To test the hypothesis that GABA would be differentially affected in the left and right nuclei during adaptation, we then performed a two-way RM-ANOVA with within-subject factors of hemisphere (Left, Right) and timepoint (Early, Late, After). In line with our hypothesis, we observed significant GABA:tCr modulation in left compared to right nuclei (significant hemisphere*time interaction: F(2,32) = 4.490, p = 0.014, partial η2 = .236; main effect of timepoint: F(2,32) = 0.054, p = 0.948, partial η2 = 0.003; Figure 3C).

Finally, to test the neurochemical specificity of the observed GABA:tCr changes in the cerebellar nuclei, we performed an equivalent RM-ANOVA to investigate adaptation-related changes in Glutamate:tCr. There was no main effect of timepoint on Glutamate:tCr, and no hemisphere*time interaction (main effect of timepoint: F(2,32) = 0.054, p = 0.948, partial η2 = 0.003, F(2,32) = 4.490, p = 0.014, partial η2 = 0.236; Figure 3F), suggesting that these findings were specific to GABA and not a reflection of more general processes.

### Adaptation-driven early GABA change correlates with adaptation

Prior studies have shown that early change in motor cortical GABA during learning of a motor sequence correlates with subsequent learning-related changes in reaction times on the task (Kolasinski, 2019; Floyer-Lea et al., 2016). We therefore hypothesised that early change in the right cerebellar nuclei GABA:tCr would correlate with error during adaptation. We observed a significant correlation between change in GABA:tCr in the right nuclei and adaptation error (Figure 4B; r = 0.64 p = 0.006), such that participants showing an early decrease in right nuclei GABA:tCr adapted their right-hand movements better to the rotation. This relationship was specific both neurochemically (change in Glutamate:tCr and adaptation r = −0.10 p = 0.693; zdiff = 2.27, p = 0.011; Figure 4C) and anatomically (change in left nuclei GABA:tCr and adaptation r = −0.16, 0.537; zdiff = 2.43, p < 0.01). To test whether our relationship between GABA:tCr and adaptation was specific to that behaviour, we tested for a relationship between GABA:tCr change and retention error, which demonstrated a significant correlation (r = −0.49 p = 0.0455). However, as expected, our metrics of adaptation and retention are highly correlated (R = −0.8185 p < 0.0001). We therefore calculated partial correlations between early GABA:tCr change and the two behavioural metrics. GABA:tCr change and adaptation were still significantly correlated when controlling for retention (r = 0.47, p = 0.03 [one-tailed]), but GABA:tCr change did not correlate with retention when controlling for adaptation (R = 0.07, p = 0.39).

**Fig. 4.**
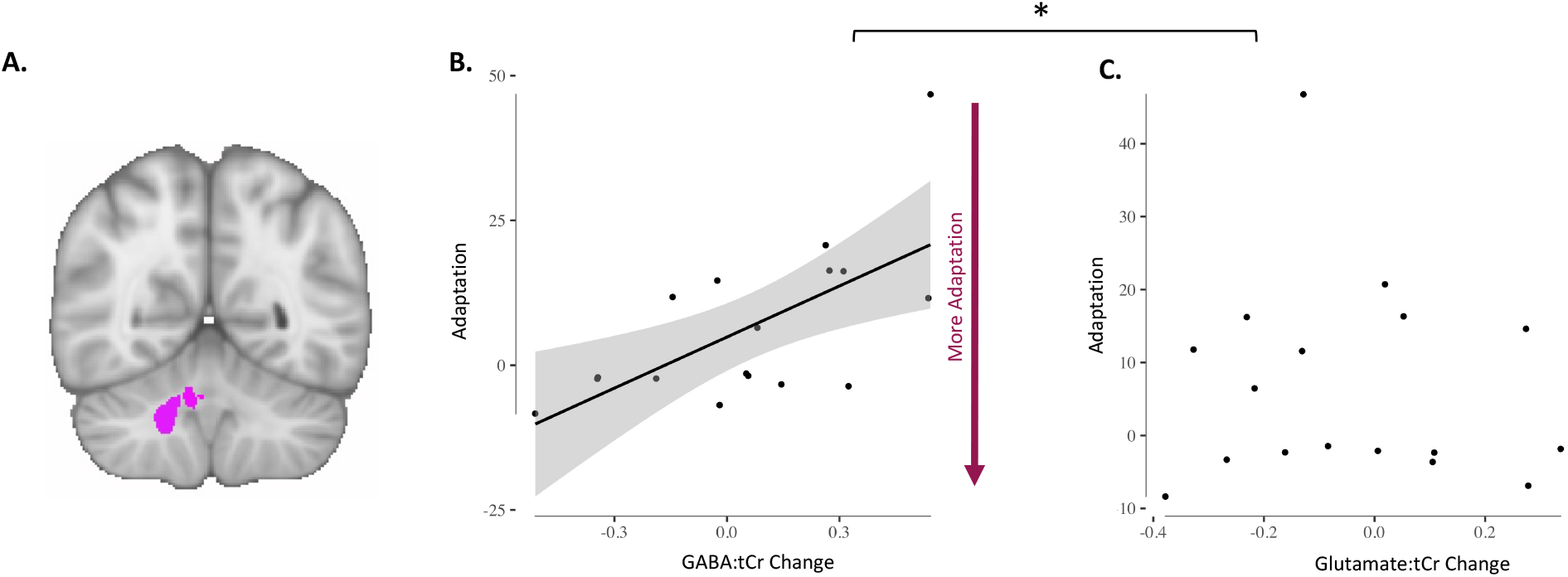
Adaptation-driven early GABA change correlates with adaptation. **A.** Right cerebellar nuclei ROI. **B.** GABA change in right cerebellar nuclei correlates with adaptation (r = 0.64 p = 0.006). Each dot is one participant. Shading indicates 95% confidence bounds of the relationship. **C.** Glutamate change in right cerebellar nuclei does not correlate with adaptation (r = −0.10 p = 0.693). Each dot is one participant. Line of best fit is not plotted for this correlation, because the correlation is not significant. * Indicates that the relationships were significantly different (zdiff (GABA change vs Glutamate change relationship) 2.27, p = 0.011).

### Adaptation-driven GABA changes correlate with cerebellar connectivity change

Finally, we wanted to determine whether our behaviour-related changes in GABA correlated with changes in cerebellar functional connectivity. We did not demonstrate a significant group mean change in cerebellar network strength after adaptation (all p > 0.05). However, there was a significant correlation between early change in right nuclei GABA:tCr and the change in cerebellar functional connectivity (r = −0.4939 p = 0.0439; Figure 5D). This relationship was both neurochemically (Glutamate change and cerebellar connectivity change: r = 0.09 p = 0.71; zdiff = −1.69 p = 0.0459; Figure 5E) and anatomically specific (GABA change and connectivity change in DMN, r = 0.2707 p = 0.29; zdiff = −2.17 p = 0.015; Figure 5C). There was no relationship between cerebellar connectivity change and adaptation error (r = −0.11 p = 0.67).

**Fig. 5.**
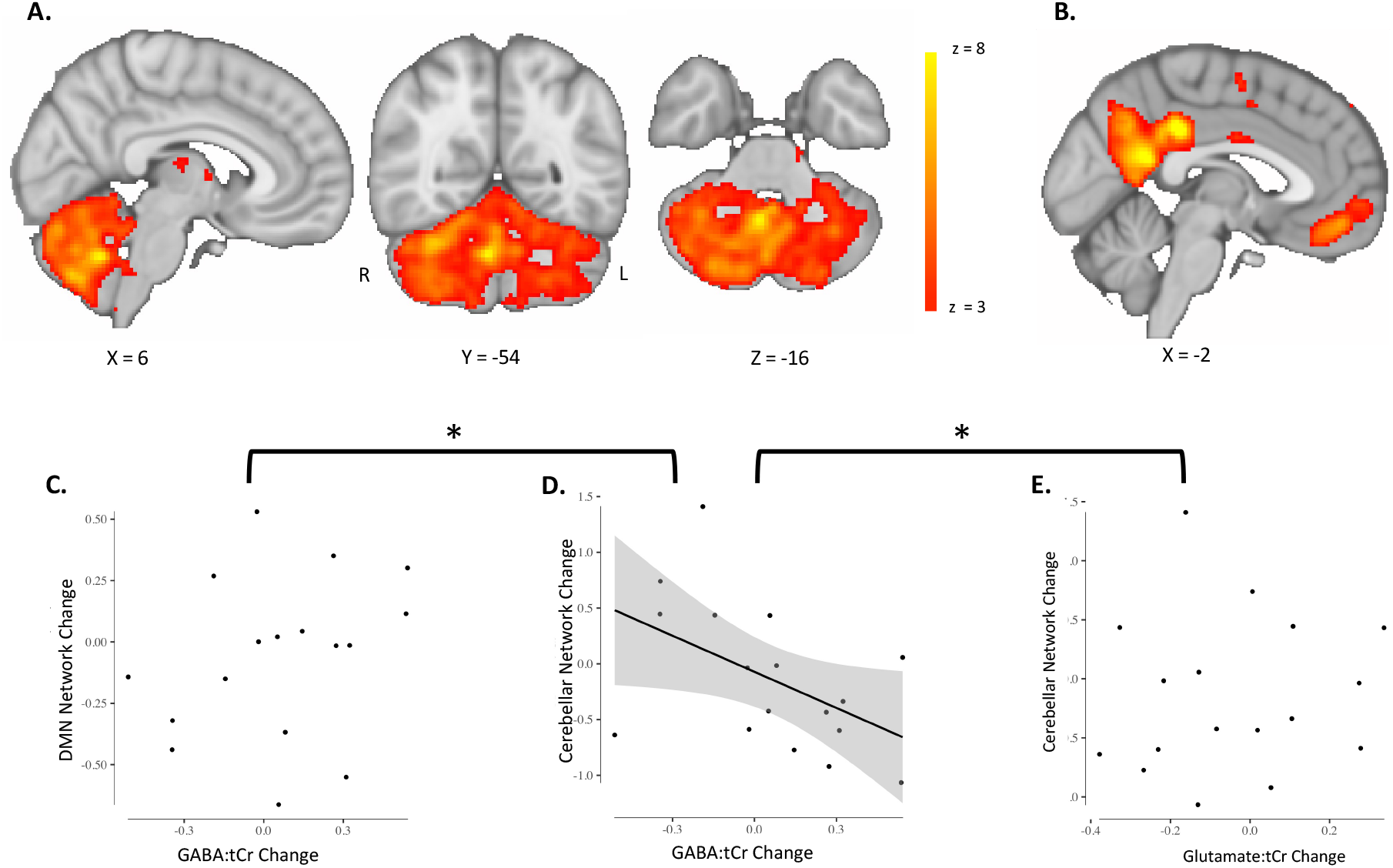
Adaptation-driven GABA changes correlate with cerebellar connectivity change. **A.** Cerebellar network. Cerebellar network that has previously been found to increase in connectivity after adaptation is shown in red-yellow. Colour bar indicates range of z-values in voxels. Brain slices are shown according to radiology convention, so left hemisphere is shown on the right (indicated by R and L surrounding the coronal slice). **B.** Default Mode Network. The Default Mode Network (DMN) was chosen as a control network. **C.** Change in Default Mode Network strength does not correlate with GABA change in right cerebellar nuclei (, r = 0.2707 p = 0.29). Datapoints are individual participants. Line of best fit is not plotted for this correlation, because the correlation is not significant. **D.** Change in cerebellar network strength correlates with GABA change in right cerebellar nuclei. Participants who show a greater decrease in early GABA in the right cerebellar nuclei also show a larger increase in cerebellar network strength (r = −0.4939 p = 0.0439). Cerebellar network strength decreases in participants who show an increase in GABA in the right cerebellar nuclei. Datapoints are individual participants. Plot shows line of best fit for correlation and 95% confidence interval. **C.** Change in cerebellar network strength does not correlate with Glutamate change in right cerebellar nuclei (r = 0.09 p = 0.71). Datapoints are individual participants. Line of best fit is not plotted for this correlation, because the correlation is not significant. * Indicates significant difference between correlations shown in C and in D as well as D and E: The correlation coefficient of Cerebellar network change and GABA change was significantly different from the correlation coefficient of DMN change and GABA change (zdiff = −2.17 p = 0.015). The correlation coefficient of Cerebellar network change and GABA change was also significantly different from the correlation coefficient of Cerebellar network change and Glutamate change (zdiff = −1.69 p = 0.0459)

## Discussion

In this study, we used a novel, spatially-resolved MRSI technique to quantify cerebellar GABA to investigate neurochemical changes during visuomotor adaptation. We found that simple movement of the right hand increases GABA in the right cerebellar nuclei and decreases GABA in the left. When controlling for these movement-related GABA changes, in line with our hypotheses we found an increase in GABA in the left cerebellar nuclei and a decrease in GABA in the right cerebellar nuclei during adaptation. This early adaptation-driven change in right cerebellar nuclei GABA correlates with performance in the right-hand adaptation task such that participants who a show greater GABA decrease adapt better, suggesting this early GABA change is behaviourally relevant. Finally, while we did not demonstrate a group mean change in cerebellar connectivity, the adaptation-driven right cerebellar nuclei GABA decrease correlated with functional connectivity changes in a network that has previously been shown to increase in strength in response to adaptation (Albert et al., 2009; Nettekoven et al., 2020). Participants who show a greater early decrease in GABA in the right cerebellar nuclei also show a greater increase in cerebellar network connectivity. These findings are anatomically specific and neurochemically specific: control analyses show no adaptation-related changes in the cerebellar cortex and no concentration changes in glutamate. Our results are also behaviourally specific and network-specific: The relationship between adaptation and right nuclear GABA change holds when controlling for retention, but there is no correlation between retention and right nuclear GABA change controlling for adaptation, suggesting this relationship is specific to adaptation. Further, right nuclear GABA change correlates with strength change in a cerebellar network, but not with strength change in the Default Mode Network, confirming the specificity of our results for functional connectivity of the cerebellum.

### Movement-related increase in GABA in ipsilateral nuclei

When groups of parallel fibres activate and cause high enough input to Purkinje cells so that Purkinje cells fire simple spikes, this results in GABA release at the cerebellar nuclei. Simple spike responses can occur in response to a large variety of vestibular and motor signals *in vivo* (Barmack and Yakhnitsa, 2013; Dean et al., 2010). For example, increases in simple spike firing have been shown during eye movements in the direction of their preferred response (Medina and Lisberger, 2008; Yang and Lisberger, 2014). There may also be population coding of hand movements within the CS as has been described for eye movements (Thier et al., 2000; Popa et al., 2019).

The increase in GABA in the right compared to left cerebellar nuclei during simple movement execution might reflect increased simple spike firing of Purkinje cells, leading to higher GABA concentration in the deep cerebellar nuclei. In the control condition participants performed simple ballistic wrist movements with veridical visual feedback, and participants therefore made no systematic movement errors. In response to these simple movements, converging parallel fibre activity may have caused Purkinje cells in the right cerebellum to fire simple spikes and, due to the absence of movement errors in the control condition, a lack of complex spikes may have increased simple spike activity of the same Purkinje cells.

### Adaptation-related decrease in GABA in ipsilateral nuclei

Unexpected target movements that result in a visual error have been shown to lead to complex spike firing on some trials (Medina and Lisberger, 2008; Yang and Lisberger, 2014). After trials where a complex spike occurred, reduced simple spike firing was found on the next trial, conversely if no complex spike occurred, simple spike firing showed a slight increase on the next trial (Yang and Lisberger, 2014). Similar decreases in simple spike firing rate after a complex spike have been observed following saccades, with a less pronounced increase in simple spikes when no complex spike was present on the previous trial (Herzfeld et al., 2018).

The systematic movement errors that occurred in our adaptation condition might therefore be expected to result in reduced simple spike firing after the complex spike signal transmitted the error information, thereby reducing inhibitory tone (GABA) in the nuclei. Consistent with this, complex spike firing has been shown to serve as a “teaching signal” (Ma et al., 2016), leading to long-term depression (LTD) of parallel fibre-Purkinje cell synapses of those parallel fibres active simultaneously with the climbing fibre input (Ito and Kano, 1982; Albus, 1971; Marr, 1969). LTD in turn has been shown to result in decreased simple spike firing (Ito and Kano, 1982; Gao et al., 2003; Yang and Lisberger, 2014) and improve motor performance (Medina and Lisberger, 2008). Hence it is plausible that a reduction in GABA concentration at the deep cerebellar nuclei could be tied to successful adaptation.

In line with the hypothesis that these adaption-related changes in GABA concentration are related to cerebellar plasticity, we demonstrated a correlation between the extent of early GABA changes in the right nuclei and adaptation. A greater decrease in GABA was related to better adaptation.

### Cerebellar connectivity change is correlated with GABA in the nuclei

The adaptation-related GABA change in the right cerebellar nuclei was correlated with functional connectivity change in a cerebellar network, such that a decrease in GABA was associated with stronger cerebellar network connectivity. This cerebellar network has previously been shown to increase in strength after adaptation, compared to after a control task (Albert et al., 2009; Nettekoven et al., 2020), in a performance-relevant manner (Nettekoven et al., 2020). We did not see a similar change here, perhaps because the restingstate scan in this study was not acquired immediately after the rotation blocks, but after interspersed open loop blocks, which may have diluted a strength increase of the cerebellar network after adaptation.

Previous work from our lab and others has shown that GABA measured from a key node of a functional network is related to functional connectivity within that network (Stagg et al., 2014; Bachtiar et al., 2015; Kapogiannis et al., 2013). This finding therefore suggests that the relationship between local inhibition and connectivity may be a general property of functional networks, even in the context of the highly different physiology and anatomy of the cerebellar and cerebral cortices.

### No GABA changes in the cerebellar cortex

In the cerebellar cortex, GABA is released by inhibitory interneurons in the molecular layer and the granule layer. Molecular layer interneurons, called basket and stellate cells, receive input from parallel fibres and inhibit Purkinje cells, resulting in di-synaptic inhibition in parallel with the monosynaptic excitatory input from granule cells to Purkinje cells (Rieubland et al., 2014; Arlt et al., 2020). In the granule layer, Golgi cells receive input from mossy fibres as well as granule cells and inhibit granule cells and other Golgi cells (Hull and Regehr, 2012). Although there is evidence that plasticity at molecular layer interneurons could regulate plasticity of parallel fibre-Purkinje cell synapses and influence adaptation in vivo (Rowan et al., 2018), there is no consensus yet on how precisely inhibitory interneurons in the cerebellar cortex contribute to adaptation learning.

We did not see significant changes in GABA in the cerebellar cortex during simple movement execution or during adaptation. It is possible that GABA concentration changes in the cerebellar cortex were not detectable with MRSI or that the placement of the MRSI grid did not optimally cover the right-hand representation in the cerebellar cortex spanning right lobules 5 and 6 in the anterior cerebellum (Diedrichsen and Zotow, 2015). Future studies should probe the role of GABA in the cerebellar cortex directly by optimizing voxel size and placement such that potential changes in GABA in the right-hand representation in the cerebellar cortex are captured.

### Conclusions

In this study, we present (to the best of our knowledge) the first investigation of cerebellar GABA changes during human visuomotor adaptation. We found movement-related GABA changes in the left and right cerebellar nuclei. When we controlled for these, we isolated GABA changes in the nuclei that were driven by adaptation. We found relationships between adaptation-driven GABA change in the right cerebellar nuclei and adaptation performance, and between adaptation-driven GABA change in the right cerebellar nuclei and change in cerebellar network connectivity. These results show the first evidence for plastic changes in cerebellar neurochemistry during a cerebellar learning task and provide the basis for developing therapeutic interventions that facilitate these naturally occurring changes to amplify cerebellar-dependent learning.

## Acknowledgments

The research was supported by the National Institute for Health Research (NIHR) Oxford Biomedical Research Centre and the NIHR Oxford Health Biomedical Research Centre. CJS holds a Sir Henry Dale Fellowship, funded by the Wellcome Trust and the Royal Society (102584/Z/13/Z). HJB holds a Wellcome Principal Research Fellowship (110027/Z/15/Z). The Wellcome Centre for Integrative Neuroimaging is supported by core funding from the Wellcome Trust (203139/Z/16/Z). For the purpose of open access, the author has applied a CC BY public copyright licence to any Author Accepted Manuscript version arising from this submission.

## Data Availability

MRI data, MRSI data and behavioural will be shared on the data sharing platform of the Wellcome Centre for Integrative Neuroimaging which is currently under development. In the interim, data is available upon request.

## Author Contributions

CRN: Designed research, performed research, analyzed data, wrote the first draft of the paper and edited the paper

LM: Performed research, analyzed data, edited the paper

WTC: Contributed analytic tools and edited the paper

UE: Contributed analytic tools and edited the paper

JC: Performed the research and edited the paper

HJB: Edited the paper

NJ: Designed the research and edited the paper

CJS: Designed research, performed research, wrote the paper and edited the paper

